# Mitochondrial uncoupler and retinoic acid synergistically induce differentiation and inhibit proliferation in neuroblastoma

**DOI:** 10.1101/2024.01.22.576741

**Authors:** Haowen Jiang, Sarah Jane Tiche, Clifford JiaJun He, Mohamed Jedoui, Balint Forgo, Meng Zhao, Bo He, Yang Li, Albert M. Li, Anh T. Truong, Jestine Ho, Cathyrin Simmermaker, Yanan Yang, Meng-Ning Zhou, Zhen Hu, Daniel J. Cuthbertson, Katrin J. Svensson, Florette K. Hazard, Hiroyuki Shimada, Bill Chiu, Jiangbin Ye

## Abstract

Neuroblastoma is a leading cause of death in childhood cancer cases. Unlike adult malignancies, which typically develop from aged cells through accumulated damage and mutagenesis, neuroblastoma originates from neural crest cells with disrupted differentiation. This distinct feature provides novel therapeutic opportunities beyond conventional cytotoxic methods. Previously, we reported that the mitochondrial uncoupler NEN (niclosamide ethanolamine) activated mitochondria respiration to reprogram the epigenome, promoting neuronal differentiation. In the current study, we further combine NEN with retinoic acid (RA) to promote neural differentiation both *in vitro* and *in vivo*. The treatment increased the expression of RA signaling and neuron differentiation-related genes, resulting in a global shift in the transcriptome towards a more favorable prognosis. Overall, these results suggest that the combination of a mitochondrial uncoupler and the differentiation agent RA is a promising therapeutic strategy for neuroblastoma.

## Introduction

Neuroblastoma, the predominant tumor in the sympathetic nervous system (97%), is the most common malignancy in infancy, diagnosed at a median age of 17 months [1]. Though the children with low-risk or intermediate-risk neuroblastoma exhibit a 5-year survival rate ranging between 90% and 95%, those with high-risk neuroblastoma face a notably lower survival rate of approximately 50% [2]. Collectively, neuroblastoma, spanning low to high risk, contributes to 15% of pediatric cancer-related fatalities. This highlights the critical need for innovations in diagnostics and therapeutics, specifically tailored to address the unique challenges posed by varying risk categories, particularly the high-risk subgroup.

Unlike adult cancers, which are predominantly epithelial and typically arise due to aging and/or exposure to mutagens, pediatric cancers are rarely epithelial. Instead, they mainly originate from mesodermal and ectodermal lineages [3]. A key aspect of pediatric neoplasms is their developmental origin, often presenting as a maturation block—a potential target for innovative treatments [3]. The neuroblastoma arises from malfunctioning differentiation of neural crest-derived cells in the peripheral sympathetic nervous system [4, 5]. Thus, promoting neuronal differentiation emerges as a therapeutic avenue for neuroblastoma. Retinoic acid (RA) has proven to be a potent inducer of neuroblastoma differentiation, exhibiting significant reductions in proliferation and increased neurite formation across various cell lines *in vitro*. The RA derivative, 13-cis RA, rapidly advanced into neuroblastoma clinical trials and is now a component of standard care [6]. Although it was shown that RA treatment increase event-free survival [7], its efficacy is constrained by numerous factors. For instance, differentiation of neuroblastoma induced by 13-cis-RA is impeded under hypoxic conditions due to histone hypo-acetylation, suppressing RA signaling gene expression [8]. In addition, it was reported DNA methyltransferase inhibitor can reduce RA-resistance in neuroblastoma and promote differentiation [9, 10], suggesting epigenetic remodeling is one major cause of RA resistance.

Our recent studies have shown that treatment with the mitochondrial uncoupler niclosamide ethanolamine (NEN) activates mitochondrial respiration and promotes neural differentiation in neuroblastoma cells [11]. Mechanistically, this treatment elevates NAD^+^/NADH, pyruvate/lactate, and α-ketoglutarate (α-KG)/2-hydroxyglutarate (2-HG) ratios, leading to promoter CpG island demethylation and epigenetic remodeling. Furthermore, it upregulates p53 and downregulates N-Myc and β-catenin signaling pathways. Even under hypoxic conditions, NEN effectively inhibits 2-HG production and suppresses hypoxia-inducible factor (HIF) signaling. When supplemented in the diet, NEN decreases tumor growth, 2-HG levels, and N-Myc/β-catenin expression in an orthotopic neuroblastoma mouse model. An integrative analysis of multiple neuroblastoma patient datasets has revealed that NEN upregulates genes associated with favorable prognosis and downregulates genes associated with unfavorable outcomes [11]. Given the differentiation-inducing effects of both retinoic acid (RA) and mitochondrial uncouplers in neuroblastoma, our research aims to determine whether combining RA with an uncoupler could synergistically enhance therapeutic differentiation effects in neuroblastoma.

In the current study, we showed that the RA+NEN combination induced proliferation arrest and neuronal differentiation in neuroblastoma synergistically. Consistently, a gene signature associated with neuronal differentiation and favorable prognosis in neuroblastoma is upregulated by the combination of RA and NEN treatment. Dietary supplementation with RA+NEN induced neuronal differentiation in an orthotopic neuroblastoma mouse model. Intriguingly, while RA treatment alone did not induce a significant shift of global transcriptome toward favorable prognosis, the RA+NEN combination was able to do so. The findings strongly indicate that employing a synergistic approach involving an uncoupler and the differentiation agent RA holds promise as an effective and specific therapeutic strategy for neuroblastoma.

## Materials and methods

### Cell culture and reagents

*MYCN*-amplified neuroblastoma cell lines SK-N-BE(2), NB16 and CHP134 were obtained from Dr. John M. Maris’ laboratory (Children’s Hospital of Philadelphia). Certificate of analysis is available from Dr. Maris’ group. All cell lines were tested and found negative for mycoplasma (MycoAlert Mycoplasma Detection Kit; Lonza). The authors did not authenticate these cell lines. All cell lines were cultured in DMEM/F12 medium (Caisson Labs, DFL15) supplemented with 1% Penicillin-Streptomycin (Gibco, 15140122), 10% FBS (Sigma, F0926, lot 17J121) and extra 1mM glutamine (Gibco, 25030081). Niclosamide ethanolamine (NEN, #17118) and 13-cis-retinoic acid (RA, # 21648) were purchased from Cayman. Anti-β-tubulin III antibody (#5568S) was purchased from Cell Signaling, and Goat anti-Rabbit IgG (H+L) Cross-Adsorbed Secondary Antibody Alexa Fluor 594 (#A-11012) was purchased from Invitrogen.

### Cell proliferation assay

The neuroblastoma cells were plated into a black 96-well plate (Corning, #3603) for treatment or pretreatment indications. After 48 or 96 hours, Hoechst 33342 Solution (20 mM, Thermo, #62249) was added to the well to achieve a final concentration of 2 µM, followed by a 5-10 minute incubation protected from light. Cell counting was conducted using an Agilent BioTek Cytation 5 (Agilent Technologies) imaging system with Gen5 software (Agilent Technologies) following the manufacturer’s instructions [12].

### Chou-Talalay analysis

The cells were seeded in a black 96-well plate (Corning, #3603) and treated to a two-fold serial dilution set-up. This set-up included an untreated control, as well as 0.25*IC50, 0.5*IC50, IC50, 2*IC50, and 4*IC50 concentrations of either RA or NEN, or a combination of both. After a 72-hour incubation period, cell counts were determined using the method described above. The data obtained from individual treatments normalized to the values obtained from untreated controls. The nature of each drug combination was assessed using a combination index (CI) according to Chou-Talalay method [13], and CI values were calculated using CompuSyn software.

### RNA isolation, reverse transcription, and real-time PCR

The procedure was performed as described before [14]. Briefly, total RNA was isolated from 60mm tissue culture plates according to the TRIzol Reagent (Invitrogen, 15596026) protocol. 3 μg of total RNA was used in the reverse transcription reaction using the iScript cDNA synthesis kit (Bio-Rad). Quantitative PCR amplification was performed on the Prism 7900 Sequence Detection System (Applied Biosystems) using Taqman Gene Expression Assays (Applied Biosystems). Gene expression levels were normalized to 18S rRNA. Data is presented as mean ± SD of three PCR reactions, and representative of three independent experiments is shown.

### RNA-seq and bioinformatics analysis

The total RNA from treatment groups (Ctrl and NEN treated, n = 3) were extracted using Trizol reagent according to the manufacturer’s instructions. The library was constructed and subjected to 150 bp paired-end sequencing on an Illumina sequencing platform (Novogene). RNA-seq analysis was performed using the kallisto and sleuth analytical pipeline. In brief, a transcript index was generated with reference to Ensembl version 67 for hg19. Paired-end mRNA-seq reads were pseudo-aligned using kallisto (v0.42.4) with respect to this transcript index using 100 bootstraps (-b 100) to estimate the variance of estimated transcript abundances. Transcript-level estimates were aggregated to transcripts per million (TPM) estimates for each gene, with gene names assigned from Ensembl using biomaRt. Differential gene expression analysis was performed using the sleuth R package across pairwise groups using Wald tests, with significant hits called with a sleuth q-value <0.05 and log2(fold change) > 0.693 or <-0.693. Gene set enrichment analysis (GSEA) was used to examine the significantly enriched pathways by comparing the normalized data of the entire RNA seq TPM dataset between groups as indicated to Molecular Signatures Database (MSigDB v7.4). All the annotated transcripts (∼21,427 features in total) with expression values were uploaded to a locally-installed GSEA tool (version 4.1.0) and compared against the Hallmark gene sets (50 gene sets). To generate the favorable or unfavorable prognosis gene sets, Kaplan-Meier assay was employed for all the genes independently by using all 11 available neuroblastoma patient databases from R2 (https://hgserver1.amc.nl/cgi-bin/r2/main.cgi). Each gene that passes filter of Kaplan-Meier assay showed significant correlation with prognosis (p-value<0.05) in relevant databases.

### Neurite outgrowth assay

The quantification of neurite was performed as described before [8]. Shortly, 1 × 10^4^ SK-N-BE(2) and NB16 cells were plated in a 12 well-plate per well. After overnight incubation, cells were treated with various conditions as indicated for 72h. Then, images were taken by a Leica Florescent Microscope DMi8 in phase contrast mode (20× magnification). The lengths of the neurites were traced and quantified using the ImageJ plugin NeuronJ[15]. For each sample, total neurite length was measured and normalized to the number of cell bodies. Data are presented as mean ± SD of three biological repeats.

### Immunofluorescence staining

SK-N-BE(2) and NB16 cells were seeded into 8-champer slides (Thermo Fisher Scientific, 154534) with a density of 6×10^3^ cells/well overnight and treated with indicated treatment for 72h. The procedure was performed as described before [8]. Cells were fixed with 4% PFA in PBS with 0.1% Tween-20 at RT for 30 minutes followed by permeabilization with 0.1%Triton X-100 in PBS at room temperature for 10 min. The cells were washed with PBS twice and blocked with 2.5% horse serum in PBS at RT for 1 h. Then, cells were subjected to immunofluorescence staining with primary antibody at 4 °C overnight. After two washes with PBS, cells were incubated with Alexa Fluor 594-conjugated anti-rabbit secondary antibody (Life Technologies) at RT for 1 h followed by staining with DAPI (Vector Laboratories, H-1800-2). Images were acquired with Leica DMi8 microscope. Data is the representative of two independent experiments.

### Mouse orthotopic neuroblastoma model

All mouse procedures were performed in accordance with Stanford University Administrative Panel on Laboratory Animal Care recommendations for the care and use of animals and were maintained and handled under protocols approved by the Institutional Animal Care and Use Committee. All procedures were performed with 1:1 gender ratio nude mice (Taconic, Hudson, NY, USA) at 7 weeks of age. Procedures and ultrasound measurements (see below) were performed under general anesthesia using isoflurane inhalation. Orthotopic tumors within the adrenal gland were created as described before [16]. Briefly, a transverse incision was made on the left flank to locate the left adrenal gland, and 1μL of phosphate buffered saline (PBS) containing 10^5^ SK-N-BE(2) or NB16 cells were injected into the adrenal gland using a 30G needle. Fascia and skin were closed in separate layers. Tumor formation was monitored by non-invasive ultrasound measurements and diet intervention were given to mice once the tumor volume reached between 5-10 mm^3^. Control diet (#D11112201) and diet with 2000ppm NEN (#D20101601) were purchased from Research Diets, Inc. The animals were euthanized when the tumor volume exceeded 1,500 mm^3^.

### Monitoring tumor growth with high frequency ultrasound

After securing the mouse in a prone position, a Visual Sonics Vevo 2100 sonographic probe (Toronto, Ontario, Canada) was applied to the left flank to locate the left adrenal gland and the tumor. The monitor of tumors by ultrasound was performed twice a week. Serial cross-sectional images (0.076 mm between images) were taken. The tumor volume was measured using the 3-D reconstruction tool (Vevo Software v1.6.0, Toronto, Ontario, Canada).

### Histological and Immunohistochemical examination of mouse orthotopic tumors

H&E and immunohistochemically stained sections were prepared from formalin-fixed and paraffin-embedded blocks of the mouse orthotopic tumors. As for immunostaining, unstained sections were heated for 30 minutes in Bond^TM^ Epitope Retrieval Solution 2 (No. AR9640; Leica Biosystems Newcastle LTD, Benton Ln, Newcastle Upon Tyne, UK), and incubated with anti-human N-Myc rabbit monoclonal antibody (#51705, Cell Signaling Technology, Inc., MA, USA at a dilution of 1:300) and S100 beta (EP32) Rabbit Monoclonal Primary Antibody (Sigma, #449R-17-RUO). The counter staining with hematoxylin was performed.

### Statistics

For cell proliferation and MS experiments, three biological repeats were used for data analysis. Results were represented as mean ± SD. The student’s t-test and one-way/ two-way ANOVA test was performed to determine the significance between groups (two-tailed, unequal variance).

## Results

### RA and mitochondrial uncoupler NEN synergistically induce neuronal differentiation in neuroblastoma cells

To investigate the differentiation effect induced by mitochondrial uncoupler NEN, RA or the combination of both, multiple NB cell lines were treated under indicated conditions for 72 hours (Figure 1A). RA alone induced obvious neurite growth in RA-sensitive cell lines CHP134, to a lesser extent in SK-N-BE(2), but showed no effect in RA-resistant cell line NB16. Consistent to our previous observation, NEN induced a significant neurite length increase in all three NB cell lines. Notably, NEN and RA treatment synergistically induced neuron differentiation morphology changes in neuroblastoma cells, as determined by the neurite length measurement and immunofluorescence staining for the neuron differentiation marker β-tubulin III (Figure 1A and B). To investigate whether RA and NEN treatments have synergistic effect on inhibiting proliferation, which is reversely associated with cell differentiation, we performed Chou-Talalay analysis with two-fold serial dilutions (Figure 1C) to determine the Combination Index (CI)[17]. Synergy was observed with the combination of RA and NEN at 0.25*IC50, 0.5*IC50, and IC50 concentrations (CI<1). The synergistic effect was masked at the high concentrations (CI>1). Next, we wonder whether this differentiation and cell cycle arrest effects could last after the treatments were removed. After a 4-day pretreatment, cells pretreated with RA or NEN alone still proliferate slower than control cells. This proliferation arrest effect was more pronounced when RA and NEN treatments were combined (Figure 1D).

**Figure 1.**
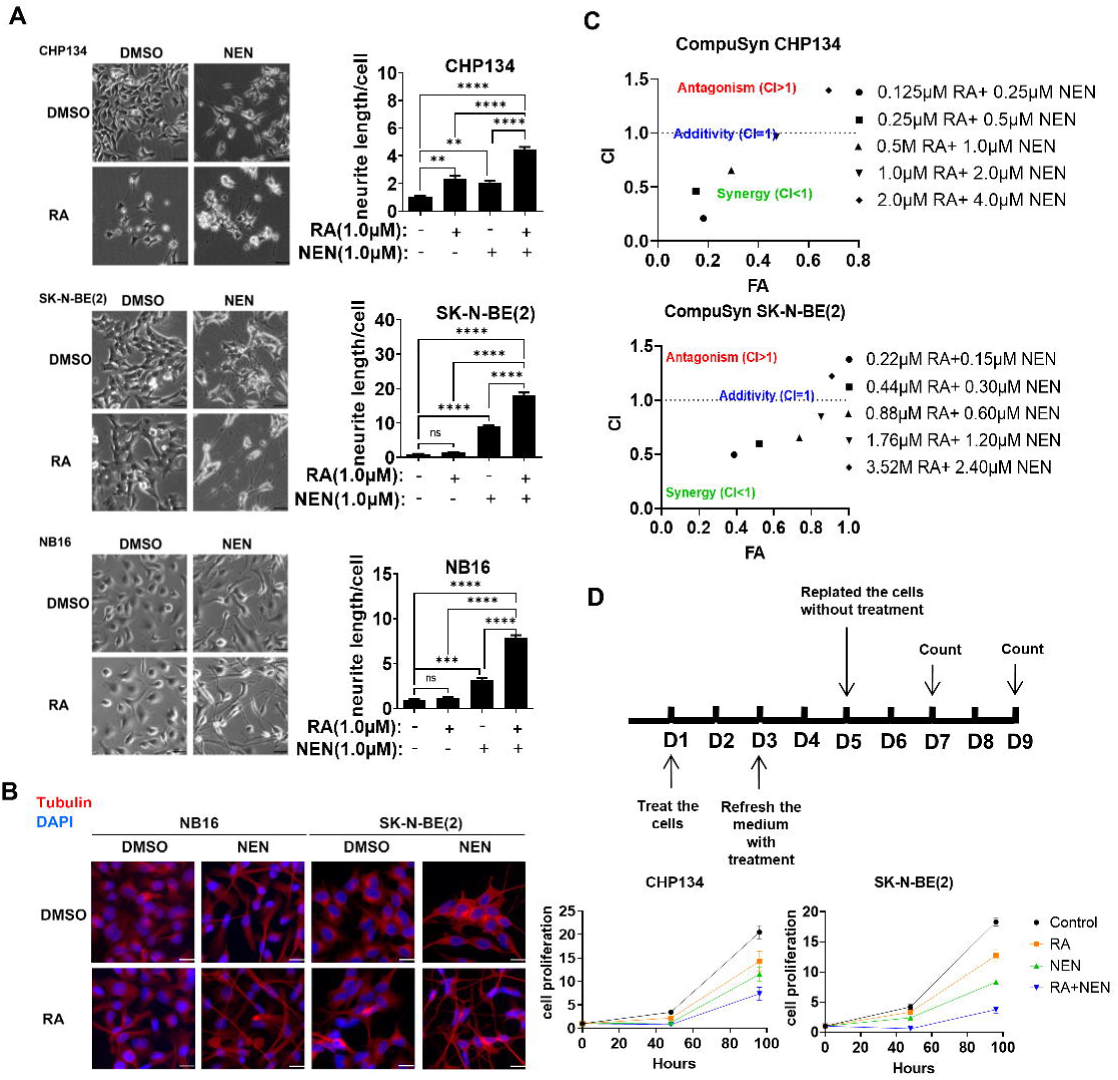
RA+NEN combination synergistically induce neuronal differentiation and proliferation arrest in neuroblastoma cells. **(A)** Left: morphological feature of neuroblastoma cells treated with indicated conditions for 96h (Scale bar: 50 μM). Right: Quantification of neurite outgrowth with NeuronJ. *P < 0.05, **P < 0.01, ***P < 0.001, ****P < 0.0001, determined by the two-tailed Student’s t-test. **(B)** Immunofluorescence staining of β-tubulin III(Red) and DAPI (Blue) in cells treated with indicated condition for 96h. Scale bar: 25μm. **(C)** Cells in a 96-well plate were treated with a series of treatments, including RA, NEN, or their combination. After 72 hours, cell counts were normalized to controls. Drug combination effects were assessed using Chou-Talalay’s method and CompuSyn software, yielding combination index (CI) values. **(D)** Cells were pretreated with RA, NEN, or their combination. After 96 hours, cell viability was determined using trypan blue, and equal numbers of viable cells from each treatment group were replated in 96 well plate. Cell counts were performed at 48 and 96 hours using Cytation 5. Cells were kept atmospherically controlled in the imaging system under the same conditions as in the incubator.

### Transcriptional profiling revealed that RA and NEN synergistically induce neuronal differentiation and activate RA-signaling pathway genes

Because cell differentiation requires transcriptome reprograming to express specific differentiation markers, we performed RNA-seq analysis on SK-N-BE(2) treated with DMSO, RA, NEN or RA+NEN for 16 hours. To understand the biological functions of these transcriptional changes (RA vs CTRL; NEN vs CTRL; RA+NEN vs CTRL), we employed Gene Set Enrichment Analysis (GSEA) using gene sets derived from Gene Ontology (GO) (Biological Process). “Proximal/distal pattern formation” and “anterior/posterior pattern specification”, two key processes during embryo development, were enriched in RA or NEN treatment alone, while the enrichment were more pronounced in the combination of RA+NEN group. Multiple neuron differentiation pathways including “neuron maturation”, “regulation of axonogenesis”, “regulation of neurogenesis”, “neural tube patterning” and “regulation of neuron projection and development” were more enriched in NEN treatment compared to RA treatment. This enrichment aligns with the observed morphological changes in differentiation. Additionally, the combination of both treatments further enhanced these pathways (Figure 2A). Conversely, both RA and NEN treatments led to a decrease in the enrichment of several pathways associated with cell proliferation. These include pathways like “DNA replication,” “positive regulation of the cell cycle process,” and “regulation of transcription involved in G1-S transition of mitotic cell cycle” (Figure 2A). The combined use of RA and NEN displayed a synergistic effect in the enrichment of these pathways, aligning with the observed synergistic impact on arresting cell proliferation (Figure 1C). In a previous publication, we identified a list of genes associated with cell differentiation and a favorable prognosis in NB based on published datasets from R2 database (Figure 2B). Intriguingly, while RA and NEN each target specific gene sets within this group, the RA+NEN combination synergistically activates the expression of all these genes. To confirm the RNA-seq results, we analyzed the expression of specific genes using RT-PCR in three neuroblastoma cell lines treated with RA, NEN, or the RA+NEN combination for 12, 24, and 48 hours (Figure 2C). Consistently, the gene expression exhibited a synergistic effect under the combined RA and NEN treatment.

**Figure 2.**
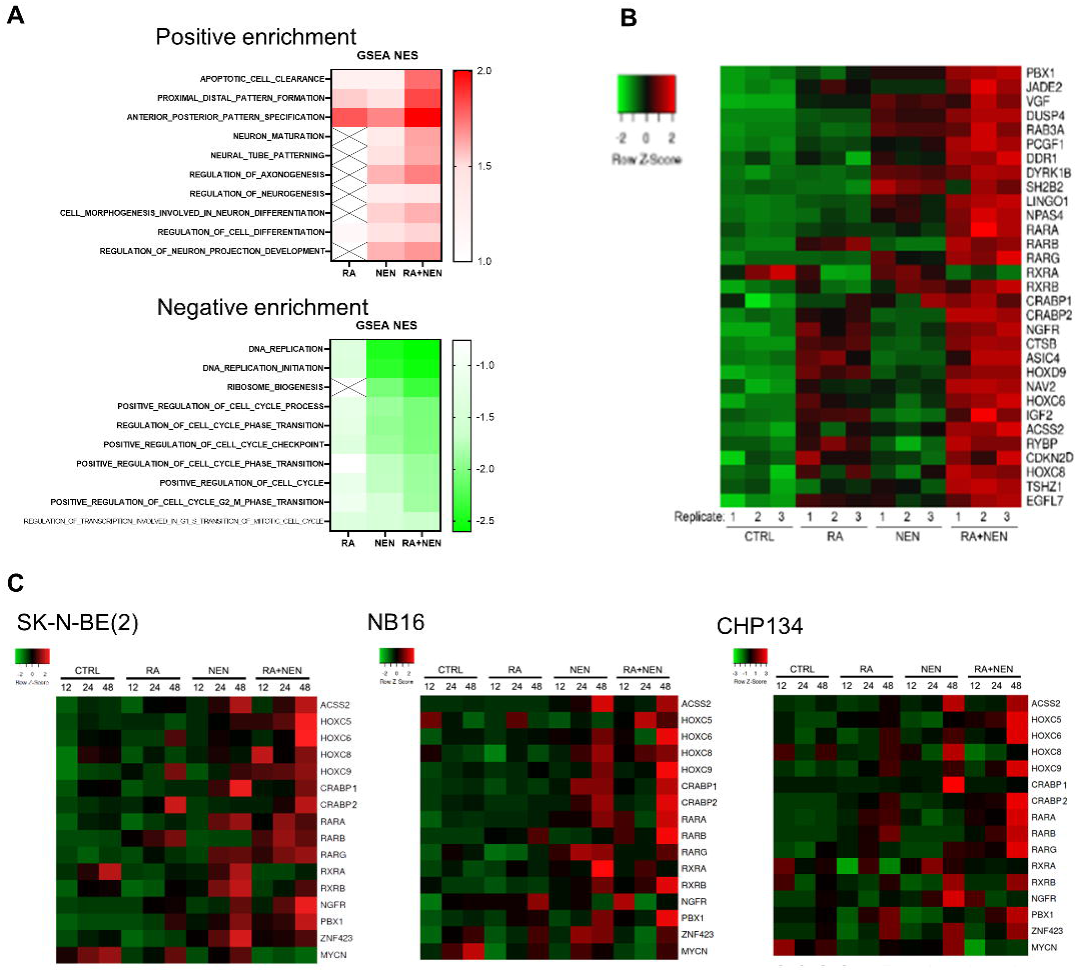
Transcriptional profiling revealed that RA and NEN synergistically induce neuronal differentiation and activate RA-signaling pathway genes. **(A)** SK-N-BE(2) cells were treated with DMSO, 1 μM RA, 1 μM NEN, and 1 μM RA and NEN for 16 hours, followed by RNA extraction and NGS. GSEA enrichment of GO Biological Process ontology was conducted on the RNA-seq experiments. **(B)** The heat map illustrates the expression of genes associated with cell differentiation and favorable prognosis from the RNA-seq experiments. **(C)** RT-qPCR analysis was performed on cell samples treated with DMSO, 1 μM RA, 1 μM NEN, and 1 μM RA and NEN for 12, 24, and 48 hours.

### RA and NEN combination drives neuronal differentiation of neuroblastoma *in vivo*

We next explored whether supplementing RA and NEN could enhance neuronal differentiation *in vivo*, using an orthotopic neuroblastoma xenograft model [12]. Once the tumors were established, the mice were administered diets containing either no supplement, 2000 ppm NEN, 83.3 ppm 13-cis-RA, or a combination of both NEN and RA. Tumor cells from the groups treated with combination of RA and NEN exhibited significantly fewer enlarged prominent nucleoli (Figure 3A and 3B), which are indicators of active ribosome biogenesis and are associated with a worse prognosis in neuroblastoma patients [18]. The control xenograft tumor displayed an undifferentiated appearance with no noticeable signs of neuroblastic differentiation. In contrast, the xenograft tumor treated with the RA+NEN combination exhibited poorly differentiated neuroblastoma subtype histology, characterized by evident neuritic process production and the formation of Homer Wright rosettes (Figure 3A). This suggests that the combined treatment effectively activates neuronal differentiation *in vivo*. Importantly, N-Myc, an oncogene commonly amplified in high-risk neuroblastoma and indicative of dedifferentiation and poor prognosis, showed significant reduction in xenografts treated with either RA or NEN. This reduction was even more pronounced in the xenografts receiving the combination treatment (Figure 3A, C). Additionally, the expression of TrkA, a marker of neuronal differentiation and favorable prognosis, increased in all treatment groups (Figure 3A, C).

**Figure 3.**
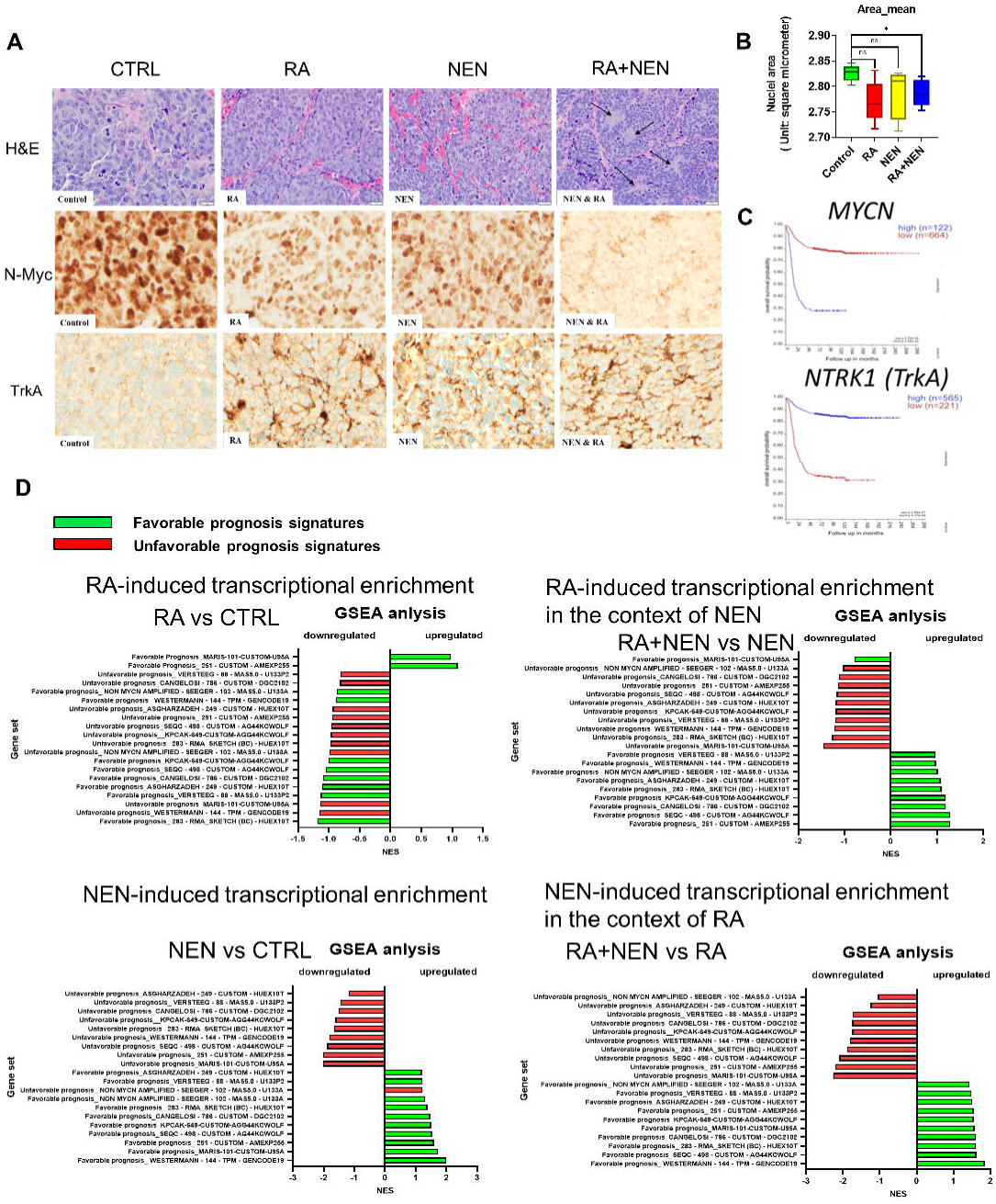
RA+NEN combination drives neuroblastoma differentiation *in vivo*. **(A)** The SK-N-BE(2) derived tumors from the CTRL, RA, NEN, and RA+NEN treatment groups were stained with Hematoxylin and Eosin (H&E) and subjected to immunohistochemical staining for N-Myc and TrkA (scale bar: 50 µm). The arrows indicate the areas of neuropil and rosette formation. **(B)** Image-based quantitation of nuclei area in **(A)**. The comparison between groups was performed using the Mann-Whitney test. **(C)** Survival curves pertaining to *MYCN* or *NTRK1* were obtained from “Tumor Neuroblastoma integrated platforms - Cangelosi - 786 - custom - dgc2102” in R2 database. **(D)** Gene Set Enrichment Analysis (GSEA) was conducted on the RNA-seq data to assess the enrichment of 10 gene sets, each indicative of either favorable or unfavorable prognosis.

To delve deeper into the clinical significance of the gene expression profile changes induced by NEN, we performed an integration analysis as described before [11]. This analysis combined the RNA-seq gene expression profiles with 10 gene sets that indicate either favorable or unfavorable prognosis. These gene sets were extracted from 10 patient studies, chosen based on the criterion that each gene’s p-value in the Kaplan-Meier analysis was less than 0.05, as sourced from the R2 database [19]. Interestingly, RA treatment, a standard component of care, upregulated genes enriched in two gene sets associated with a favorable prognosis and downregulated genes in sets linked to an unfavorable prognosis (Figure 3D). However, it is notable that RA treatment also downregulated genes in eight gene sets associated with a favorable prognosis. On the other hand, NEN treatment upregulates all the favorable prognosis gene sets and one unfavorable set. Additionally, NEN downregulates the remaining unfavorable gene sets [11]. Significantly, the transcriptional changes induced by RA+NEN combination (RA+NEN vs RA), shift towards a favorable prognosis. In this context, RA+NEN combination consistently downregulates genes across all unfavorable prognosis gene sets and upregulates genes across all the favorable ones (Figure 3D). This suggests that the combination of RA and NEN is likely to be more therapeutically effective than using RA alone.

## Discussion

Compared to adult cancer genomes, pediatric cancer genomes exhibit a significantly lower mutation rate [20]. Increasing evidence suggests that most pediatric cancers result from developmental disorders due to epigenetic dysregulation [21, 22]. Despite these fundamental differences, many pediatric cancer patients continue to receive the traditional therapies as adult patients, including surgery, radiation, and chemotherapy. Consequently, pediatric cancer survivors suffer side effects such as secondary cancer, organ toxicity, and cognitive impairments. It is essential to develop less harmful, more targeted therapeutic strategies for pediatric cancer patients.

RA-based differentiation therapy has been successfully applied in the treatment of multiple types of cancer, including acute promyelocytic leukemia (APL), teratocarcinoma, and neuroblastoma (NB)[23]. RA induces differentiation by activating the nuclear retinoic acid (RAR) and retinoid X (RXR) receptor family members. This forms RAR/RXR heterodimers that bind to retinoic acid response elements (RAREs) to activate gene expression [24], including of RARs themselves. *In vitro,* RA also has strong inductive effects on proliferation arrest, apoptosis, and differentiation in solid tumor cell lines, but the results from clinical trials are less promising [25, 26]. NB is one of the most common solid cancer types in childhood, with over 600 cases per year in the U.S. In high-risk patients, disease relapse occurs frequently after surgery or chemotherapy and eventually becomes fatal. The effect of RA has been mostly studied in NB among solid tumors because it induces clear neuronal differentiation morphology and proliferation arrest in many NB cell lines. Despite the significant response observed *in vitro*, no significant therapeutic responses were observed in clinical trials testing RA as a single agent in NB patients [25, 27, 28]. One clinical trial showed that RA significantly increased event-free survival in patients who took myeloablative therapy followed by autologous bone marrow transplantation (ABMT) or intensive chemotherapy [7]. However, even in the ABMT+RA group, over 40% of the patients did not respond to treatment and died within five years [29]. In addition, another similar study failed to show a significant response from RA treatment [30, 31]. RA-responsive NB cells can develop RA resistance, particularly under hypoxic conditions [8, 32]. These data suggest that it is necessary to develop combination therapeutic strategies to overcome RA resistance.

It was reported DNA methyltransferase inhibitor (DNMTi) can reduce RA-resistance in neuroblastoma and promote differentiation [9, 10], suggesting CpG island DNA hypermethylation is one major cause of RA resistance. Although DNMTi 5-azacytidine (5-AZA) and 5-aza-2’-deoxycytidine (decitabine) have been approved by the FDA and EMA to treat hematopoietic malignance [33], the overall efficacy in solid tumors has been less promising compared to hematological cancers [34]. There are several limitations of using DNMTi in clinic: First, they only block new methylation formation and do not remove existing methylation [35], so the ‘demethylation’ relies on cell division. Second, because DNMTi incorporate into DNA, they are considered as weak mutagens [35]. Third, they cause myelosuppression and gastrointestinal toxicities [33]. Thus, the lack of an efficient demethylation agent represents a big clinical therapeutic hurdle.

Recently, our research has led to the development of a more advanced therapeutic strategy aimed at remodeling the cancer epigenome and reducing hypermethylation of CpG island DNA through metabolic intervention [11]. According to Otto H. Warburg, the Warburg effect leads to cellular dedifferentiation, which initiates tumorigenesis [36]. Therefore, reversing the Warburg effect should reactivate the cellular differentiation program and reverse tumorigenesis [37]. We discovered that, as a mitochondrial uncoupler, NEN activates mitochondrial respiration to reverse the Warburg effect, leading to epigenetic remodeling and CpG island demethylation [11]. Surprisingly, in addition to promoter CpG demethylation, treatment with NEN also leads to an increase in gene body methylation. Notably, the methylation changes induced by NEN predominantly occur within the gene bodies of neuronal differentiation genes. This suggests that the methylation status in these regions may play a critical role in regulating gene expression and determining cell fate. Furthermore, our research revealed that mitochondrial uncoupling suppresses reductive carboxylation, a crucial metabolic pathway that contributes to cancer proliferation when the electron transport chain (ETC) is compromised [38].

The primary finding of the study demonstrates that the combination of RA and NEN synergistically induces neuronal differentiation in neuroblastoma cells, both *in vitro* and *in vivo.* The synergistic effect is observed through increased neurite length, upregulation of neuron differentiation markers, and a shift of global transcriptome toward to favorable prognosis. The combination not only induces proliferation arrest but also sustains its effects even after treatment removal, indicating a potential long-lasting impact on inhibiting cell proliferation. The study’s results highlight the molecular mechanisms behind RA and NEN-induced differentiation. Intriguingly, the combination of RA and NEN demonstrates efficacy in an orthotopic neuroblastoma xenograft model, with decreased expression of the oncogene N-Myc, and increased expression of the differentiation marker TrkA.

Because niclosamide is an FDA-approved oral antihelminth drug with very low toxicity (children can take 1g/day), it is expected that if proven effective, this metabolic treatment will be able to translate into clinical use, especially in patients with relapsed tumors. Since a mitochondrial uncoupler targets the biochemical regulation of the epigenome instead of a single mutation or altered pathway, we expect that the therapeutic strategy we establish in this study could be applied to the treatment of other types of cancer as well [39]. In several studies, niclosamide or NEN alone had anti-tumor effects in a variety of *in vitro* and preclinical tumor models [40–44]. However, the effects of using niclosamide alone in clinical trials are not promising, suggesting that combination therapy is needed. NEN has not yet been tested in clinical trials.

The study’s strengths lie in its comprehensive approach, combining in vitro experiments, transcriptomic analysis, and *in vivo* validation. The observed synergistic effects underscore the potential clinical impact of combining RA and NEN in treating high-risk neuroblastoma cases, where conventional therapies often fall short. However, some questions and considerations arise. Further investigation is warranted to elucidate the long-term safety and potential side effects of the combination therapy. Additionally, clinical trials will be crucial to validate the efficacy of this approach in human patients. The study provides a solid foundation for future research and clinical exploration, offering hope for improved outcomes in the challenging landscape of pediatric neuroblastoma treatment.

## Acknowledgement

This work was supported by a Stanford Maternal and Child Health Research Institute Research Scholar Award (2020), an American Cancer Society Research Scholar Grant (RSG-20-036-01), and an Agilent Solutions Innovation Research Award (SIRA) to J.Y. K.J.S. was supported by NIH grants R01DK125260, P30DK116074, American Heart Association 23IPA1042031, MCHRI, The SPARK Program at Stanford, the Weintz Family COVID-19 research fund, The Jacob Churg Foundation, Stanford Maternal and Child Health Research Institute, the Stanford School of Medicine, and the Stanford Cardiovascular Institute (CVI). The American Heart Association (AHA) Postdoctoral fellowship (905674) and the K99/R00 NIH NIAMS Pathway to Independence Award (K99AR081618) supported M.Z. We thank Drs. Maximilian Diehn and Beverly S. Mitchell (Stanford University) for valuable advice and suggestions.

